# An aggrecan fragment drives osteoarthritis pain through Toll-like receptor 2

**DOI:** 10.1101/160432

**Authors:** Rachel E. Miller, Richard J. Miller, Shingo Ishihara, Phuong B. Tran, Suzanne B. Golub, Karena Last, Amanda J. Fosang, Anne-Marie Malfait

## Abstract

Pain is the predominant symptom of osteoarthritis, but the connection between joint damage and the genesis of pain is not well understood. Loss of articular cartilage is a hallmark of osteoarthritis, and it occurs through enzymatic degradation of aggrecan by ADAMTS-4/5-mediated cleavage in the interglobular domain (E^373-374^ A). Further cleavage by MMPs (N^341-342^ F) releases a 32-amino-acid aggrecan fragment (32-mer). We investigated the role of this 32-mer in driving joint pain. We demonstrated that the 32-mer excites dorsal root ganglion (DRG) nociceptive neurons, both in culture and in intact explants. Treatment of cultured sensory neurons with the 32-mer induced them to express the pro-algesic chemokine, MCP-1/CCL2. These effects were mediated through Toll-like receptor (TLR)2, which we demonstrated was expressed by nociceptive neurons. In addition, intra-articular injection of the 32-mer provoked knee hyperalgesia in wild-type but not *Tlr2* null mice. Blocking the production or action of the 32-mer in transgenic mice prevented the development of knee hyperalgesia in a murine model of osteoarthritis. These findings suggest that the aggrecan 32-mer fragment directly activates TLR2 on joint nociceptors and is an important mediator of the development of osteoarthritis-associated joint pain.

## Introduction

An early event in the pathogenesis of osteoarthritis is enzymatic cleavage of the major cartilage proteoglycan, aggrecan, in the interglobular domain (E^373^ ↓^374^ A) (1-)3 by ‘a disintegrin and metalloproteinase with thrombospondin motif 4’ (ADAMTS-4) or ADAMTS-5 (4, 5). This results in loss of the bulk of the aggrecan molecule from the articular cartilage, and therefore, this cleavage step is critical for the development of osteoarthritis (6, 7). Accordingly, many pharmaceutical programs over the last two decades have focused on developing disease-modifying osteoarthritis drugs targeting ADAMTS-4 and ADAMTS-5 (8-10).

Once aggrecan is cleaved in the interglobular domain by ADAMTS-4/ADAMTS-5, the remaining 70-kDa N-terminal aggrecan fragment is retained in the cartilage matrix and is subsequently cleaved by matrix metalloproteinases (MMPs) at N^341^ ↓^342^ F, releasing a 32-amino acid fragment (32-mer, ^342^ F-E^373^) (11). This 32-mer has been identified in synovial fluid from osteoarthritis patients (12), and we have demonstrated that this fragment can promote pro-inflammatory signaling in chondrocytes, synovial fibroblasts and peritoneal macrophages, through the activation of TLR2 (13). We recently reported that nociceptors can directly respond to Damage-Associated Molecular Patterns (DAMPs) present in the osteoarthritis joint, such as S100A8 and α2-macroglobulin, by activating TLR4 receptors expressed by joint nociceptors (14). Thus, it is likely that nociceptors might also sense and respond to other DAMPs, particularly those released from the cartilage extracellular matrix as a result of ongoing degradation.

## Results and Discussion

We sought to determine whether the 32-mer aggrecan fragment produces pro-algesic effects. We first examined its ability to directly excite DRG neurons by examining its effects on intracellular calcium mobilization, [Ca^2+^] _i_. Cultured DRG neurons from wild-type mice rapidly responded to 32-mer peptide as indicated by increased [Ca^2+^]_i_ in 23% of neurons, suggesting that a subpopulation of DRG neurons express excitatory receptors for this protein fragment (Table 1, Supplementary Figure 1A). Scrambled control peptide induced responses in 6% of neurons (Table 1). All 32-mer responses were seen in small-to-medium-diameter neurons, consistent with the size of nociceptors (Supplementary Figure 1B). In addition, the majority of neurons that responded to the 32-mer peptide also responded to capsaicin (55/71 neurons = 77%), demonstrating that a subset of TRPV1 (transient receptor potential cation channel subfamily V member 1)-expressing nociceptors can respond to 32-mer. The synthetic TLR2 ligand, Pam3CSK4, also induced responses in 12% of neurons (9/76) (Supplementary Figure 1C), primarily in small-to-medium diameter, capsaicin-sensitive neurons. Finally, in DRG cultures from *Tlr2* null mice, only 5% of neurons responded to 32-mer, suggesting that excitation is dependent on this signaling pathway (Table 1). In order to show that 32-mer responses are not an artifact of cell culture, calcium imaging was also performed using intact DRG from Pirt-GCaMP3 expressing mice (14, 15). In three intact DRG explants, 6% of capsaicin-sensitive neurons (12/204) responded to 32-mer peptide, while scrambled peptide and vehicle solution each elicited responses in only 2% of capsaicin-sensitive neurons (4/204) (p=0.037) (Supplementary Figure 1D), supporting the cell culture findings.

**Table 1.** Intracellular calcium responses in cultured DRG neurons

|  | # neuronal responses / total # neurons | % responding | p-value (Chi-square test) |
| --- | --- | --- | --- |
| WT: 32-mer peptide (3 μM) | 71/309 | 23% | p<0.0001 |
| WT: Scrambled peptide (3 μM) | 18/299 | 6% | |
| *Tlr2* null: 32-mer peptide (10 μM) | 3/57 | 5% | |

In order to examine the potential pro-algesic effects of 32-mer signaling in DRG neurons, we stimulated cultured DRG cells from wild-type mice with 32-mer peptide and measured release of the chemokine, monocyte chemoattractant protein-1 (MCP-1/ CCL2) into culture medium. We previously found that MCP-1 is upregulated by DRG neurons in experimental osteoarthritis induced by surgical destabilization of the medial meniscus (DMM) and acts as a key mediator of osteoarthritis pain (16). Overnight incubation of cultured DRG cells with 3 or 30 μM 32-mer peptide resulted in significant upregulation in MCP-1 production compared with unstimulated cells (3.1-fold (3 μM) and 3.5-fold (30 μM), p<0.0001) (Figure 1A). The highest concentration of scrambled peptide (30 μM) did not induce MCP-1 production (0.8-fold, p>0.9999 *vs*. unstimulated). Since this portion of the aggrecan molecule can be modified by keratan sulfate chains (17-)20, we next tested whether the glycosylated native aggrecan 32-mer fragment can also stimulate DRG cells using native 32-mer purified from porcine cartilage. We found that native 32-mer stimulated DRG cells to produce elevated levels of MCP-1 compared with unstimulated cells (3.1-fold (10 μM), p<0.01) (Figure 1B).

**Figure 1.**
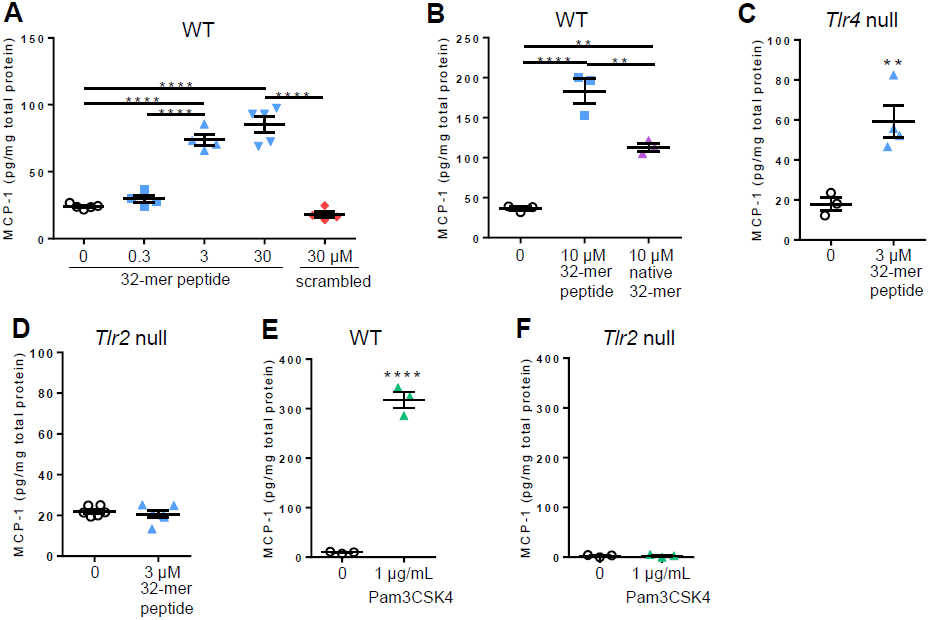
The 32-mer upregulates MCP-1 protein production through TLR2 in cultured DRG cells. **A**) Wild-type DRG cells were stimulated with 0-30 μM synthetic 32-mer or scrambled peptide. Repeated 0 vs 3 μM synthetic 32-mer in 4 independent cultures. **B**) Wild-type DRG cells were stimulated with 10 μM synthetic or native porcine 32-mer in the presence of 30 μg/mL polymyxin-B. Representative of two independent experiments. **C**) *Tlr4*^-/-^ DRG cells were stimulated with 0 vs 3 μM synthetic 32-mer. Representative of three independent experiments. **D**) *Tlr2*^-/-^ DRG cells were stimulated with 0 vs 3 μM synthetic 32-mer. Representative of three independent experiments. **E**) Wild-type DRG cells were stimulated with 0 vs 1 μg/mL synthetic TLR2 ligand Pam3CSK4. Representative of three independent experiments. **F**) *Tlr2*^-/-^ DRG cells were stimulated with 0 vs 1 μg/mL synthetic TLR2 ligand Pam3CSK4. Representative of two independent experiments. **p<0.01, ***p<0.001,****p<0.0001; mean±SEM.

Which receptors mediate the effects of the 32-mer on DRG nociceptors? In order to answer this question, we prepared DRG cultures from *Tlr2* null or *Tlr4* null mice. Stimulation with 32-mer peptide (3 μM) produced increased MCP-1 in *Tlr4* null (3.3-fold, p<0.01) (Figure 1C), but not in *Tlr2* null DRG cultures (0.9-fold, p=0.5) (Figure 1D), suggesting that these effects are mediated through the activation of TLR2. Wild-type DRG cells also responded to Pam3CSK4 (10 μM, 7.7-fold, p<0.0001) (Figure 1E) while *Tlr2* null DRG cells did not (Figure 1F), providing additional evidence that TLR2 signaling can lead to increased MCP-1 expression by these neurons. TLR2 is known to be expressed in a variety of cells as functional heterodimers with either TLR1 or TLR6, but no reports have demonstrated expression by DRG neurons (21). Therefore, we immunostained DRG sections from wild-type naïve mice and found that 17±2% of all DRG neurons stained positive for both TLR1 and TLR2, and 12±2% of all DRG neurons stained positive for both TLR6 and TLR2 (Supplementary Figure 2). In addition, naïve *Tlr2-lacZ*^*+/-*^ reporter mice also demonstrated TLR2 expression in DRG neurons (Supplementary Figure 3).

Next, we aimed to determine whether the 32-mer has the potential for generating pain *in vivo*. We first injected Pam3CSK4 into the knee cavity of naïve wild-type mice, which caused knee hyperalgesia in a dose-dependent fashion (Figure 2A). An intra-articular injection of lidocaine, administered once peak Pam3CSK4-induced knee hyperalgesia was established, reversed knee hyperalgesia, indicating a direct effect of peripheral nerves in mediating the observed hyperalgesia (Figure 2B). *Tlr2* null mice did not develop knee hyperalgesia following injection of Pam3CSK4 into the knee cavity, indicating that hyperalgesia is indeed induced through the TLR2 pathway (Figure 2C). Next, we directly tested the effects of the 32-mer by injecting either the 32-mer or scrambled peptides into the knee cavity of naïve mice. We observed that the 32-mer but not the scrambled peptide induced acute knee hyperalgesia in wild-type (Figure 2D) but not *Tlr2* null mice (Figure 2E).

**Figure 2.**
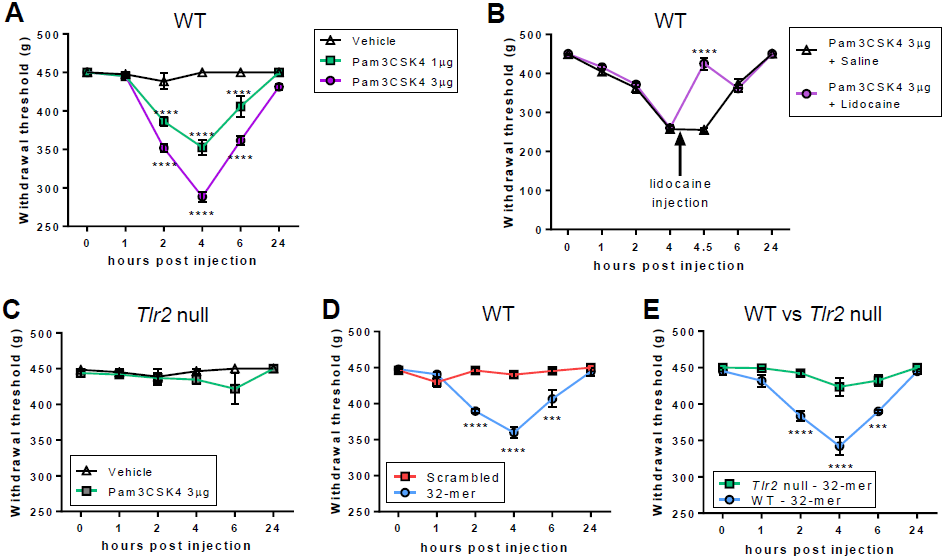
Intra-articular injection of synthetic 32-mer of Pam3CSK4 induces knee hyperalgesia in naïve mice. **A**) Intra-articular injection of vehicle, 1 or 3 μg synthetic TLR2 ligand Pam3CSK4 in wild-type mice. n=4 mice/treatment. **B**) Intra-articular injection of 3 μg synthetic TLR2 ligand Pam3CSK4 in wild-type mice followed by a second intra-articular injection of vehicle or lidocaine (20 mg/kg). n=5 mice/treatment. **C**) Intra-articular injection of vehicle or 3 μg synthetic TLR2 ligand Pam3CSK4 in *Tlr2*^*-/-*^ mice. n=4/vehicle; n=6/Pam3CSK4. **D**) Intra-articular injection of 10.5 μg synthetic 32-mer or scrambled peptide in wild-type mice. n=4/scrambled; n=5/32-mer. **E**) Intra-articular injection of 10.5 μg synthetic 32-mer peptide in wild-type or *Tlr2*^*-/-*^ mice. n=5/strain. ***p<0.001, ****p<0.0001; mean±SEM.

These data indicate that activation of TLR2 receptors by the 32-mer can elicit acute pain. In order to determine whether this process contributes to pain in active osteoarthritis, we explored the role of the 32-mer in generating knee hyperalgesia in an experimental model of osteoarthritis induced by DMM surgery. We have previously validated the DMM model as a suitable preclinical model that captures the long-term progression of osteoarthritis and associated pain (22, 23). We performed DMM surgery in wild-type, “ Chloe”, and *Tlr2* null mice. Chloe mice are a transgenic line in which the matrix metalloprotease-cleavage site (N^341^ ↓^342^F) in the aggrecan interglobular domain is eliminated by amino acid changes, thus preventing production of the 32-mer fragment (24). However, since the aggrecanase-cleavage site (E^373^ ↓^374^ A) in the interglobular domain is not modified, Chloe mice still develop osteoarthritis after DMM surgery (6). In this study, Chloe mice developed more cartilage degeneration and larger osteophytes than wild-type mice four and sixteen weeks after surgery (Supplementary Figure 4A,B). Subchondral bone sclerosis was also increased in Chloe mice compared with wild-type mice 4 weeks after DMM (Supplementary Figure 3C). Similar synovial changes were seen in both wild-type and Chloe mice throughout the 16 weeks (Supplementary Figure 4D). Wild-type and *Tlr2* null mice developed similar joint damage by 16 weeks after DMM (Supplementary Figure 4).

Similar to our previous results (25), DMM surgery caused pronounced knee hyperalgesia in wild-type mice by 2 weeks after surgery compared with sham mice, and hyperalgesia slowly resolved through 16 weeks (Figure 3A). Four weeks after DMM, intra-articular injection of lidocaine reversed knee hyperalgesia (Figure 3B), indicating that it is a locally-generated pain-related behavior. We have also previously demonstrated that knee hyperalgesia is reversed by systemic injection of morphine (25). In contrast, Chloe mice were protected from developing knee hyperalgesia until 12 weeks after DMM surgery compared with wild-type mice (Figure 3C), suggesting that the 32-mer fragment mediates the development of knee hyperalgesia. 32-mer peptide injected into the knee joint of naïve Chloe mice induced knee hyperalgesia similar to wild-type naïve mice (Figure 3D, Figure 2D), confirming that these mice retain the ability to respond to the 32-mer. *Tlr2* null mice were also protected from knee hyperalgesia up to 16 weeks after DMM surgery (Figure 3D), supporting the hypothesis that 32-mer signaling through TLR2 activation plays a key role in mediating early phase knee hyperalgesia in this model.

**Figure 3.**
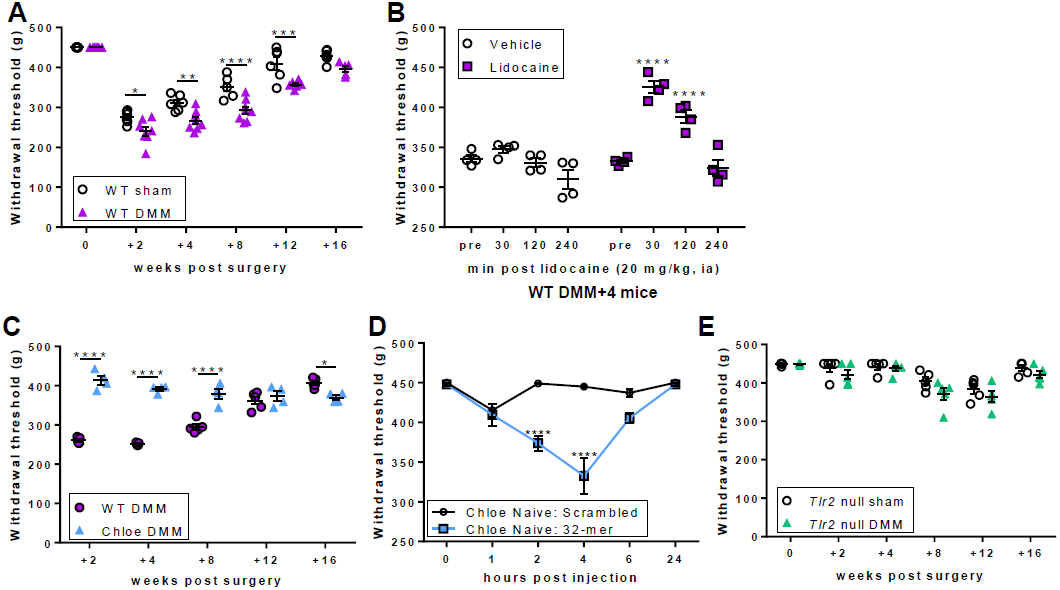
Blocking the production or action of the 32-mer protects against early knee hyperalgesia following DMM surgery. **A**) Time course of knee hyperalgesia in wild-type mice following DMM surgery. n=6-8 mice/group. **B**) Intra-articular injection of vehicle or lidocaine (20 mg/kg) four weeks after DMM surgery. n=4 mice/treatment. **C**) Time course of knee hyperalgesia in wild-type (n=5) *vs*. Chloe (n=4 mice) after DMM surgery. **D**) Intra-articular injection of 10.5 μg synthetic 32-mer or scrambled peptide in naïve Chloe mice. n=4/group. **E**) Time course of knee hyperalgesia in *Tlr2-/-* mice following sham or DMM surgery. n=5 mice/group. *p<0.05, **p<0.01, ***p<0.001,****p<0.0001; mean±SEM.

Finally, in order to test whether the 32-mer plays a role in mediating other DMM-associated chronic pain behaviors, we tested whether mechanical allodynia of the ipsilateral hindpaw was reduced after blocking the creation or action of the 32-mer. We found that wild-type, Chloe, and *Tlr2* null mice developed similar patterns of mechanical allodynia in the ipsilateral hindpaw through 16 weeks after DMM (Supplementary Figure 5), suggesting that the primary action of the 32-mer aggrecan fragment on pain is locally within the knee joint.

Toll-like receptor signaling has been implicated in a variety of neuro-immune processes (21, 26). TLR2 has been shown to be expressed by satellite glial cells in the DRG, and by microglia, astrocytes, and neurons within the central nervous system (21, 27). In nerve injury models, *Tlr2* null mice are partially protected from developing mechanical allodynia (28), microglial activation in the spinal cord (28), and macrophage infiltration into the DRG (29). Here, we demonstrate for the first time that TLR2 can be co-expressed with TLR1 and TLR6 by DRG sensory neurons, joining the list of other TLRs (TLR3, TLR4, TLR5, TLR7, and TLR9) that have also been shown to be expressed by DRG sensory neurons (14, 21, 30).

A select group of extracellular matrix molecules, including low molecular weight hyaluronan, fibronectin-EDA, fragments of fibronectin, tenascin C, and biglycan have been shown to act as DAMPs, signaling through TLR2 and TLR4 on non-neuronal cells (31). Recently, we have added the 32-mer fragment of aggrecan to this list, by demonstrating that it can upregulate catabolic signaling through TLR2 expressed by chondrocytes, synovial fibroblasts, and macrophages (13). Chondroitin-sulfate proteoglycans, including aggrecan, have been shown to play a role in guiding the development and growth of the sensory nervous system as well as regeneration following injury (32). Here, we show for the first time that an extracellular matrix fragment, specifically, the 32-mer aggrecan fragment, can signal through TLR2 on sensory neurons. It will be interesting to investigate the effects of other extracellular matrix fragments on sensory neurons in future work.

In conclusion, we have demonstrated that activation of TLR2 receptors expressed by nociceptors can promote pro-algesic signaling, particularly the production of the chemokine MCP-1 by these neurons, which we have previously identified as an important mediator of osteoarthritis pain (23). Our findings suggest that the 32-mer fragment of aggrecan directly activates TLR2 expressed by nociceptors within the knee joint, driving the development of osteoarthritis-associated knee hyperalgesia. These results provide new evidence indicating that molecules derived from the process of ongoing cartilage degradation can directly act upon nociceptors thereby integrating osteoarthritis pain with other aspects of joint degeneration.

## Methods

Additional methods are provided in the Supplemental Methods.

### Statistics

For calcium imaging experiments, Chi-squared tests were used to compare the number of responses. For MCP-1 stimulation experiments, one-way ANOVA with Bonferroni post-tests or unpaired t-tests assuming equal variances were used to compare the groups of interest. For knee hyperalgesia following intra-articular injection time courses, a repeated measures two-way ANOVA with Bonferroni post-tests was used to compare vehicle *vs*. drug at each time point. For knee hyperalgesia time courses in mice following DMM surgery, a two-way ANOVA with Bonferroni post-tests was used to compare responses at each time point. For mechanical allodynia time courses, one-way ANOVA with Bonferroni post-tests was used to compare each time point to time 0. For knee histopathology, data were analyzed using two-way ANOVA with Bonferroni post-tests or Mann-Whitney test. All analyses were carried out using GraphPad Prism version 6.07 for Windows (GraphPad Software, San Diego, CA). Results are presented as mean ± SEM or median ± IQR, as indicated.

### Study approval

All experiments were approved by the Rush University Institutional Animal Care and Use Committee.

## Author Contributions

REM, AMM, and RJM were responsible for overall study design. REM performed calcium imaging and MCP-1 experiments, collected knee joints for histopathology, and performed statistical analyses. SI performed DMM surgery and assessed knee hyperalgesia and mechanical allodynia, and other *in vivo* techniques. PBT performed the immunohistochemistry. SBG, KL and AJF supplied the synthetic 32-mer peptide, native 32-mer fragment, and Chloe mice. AJF also provided input on overall experimental design and analysis of the data. The manuscript was written by REM, RJM, and AMM. All authors were involved in the design of the parts of the study they executed, discussed the design and results, commented on the manuscript and approved the final version.

## Acknowledgements

Rachel Miller was supported by the US National Institutes of Health/National Institute of Arthritis and Musculoskeletal and Skin Diseases (NIH/NIAMS) (F32AR062927 and K01AR070328). Anne-Marie Malfait (R01AR064251 and R01AR060364) and Richard Miller (R01AR064251) were supported by NIAMS. Amanda Fosang was supported by the National Health & Medical Research Council (NHMRC) of Australia (1060222).

## Notes

The authors have declared that no conflict of interest exists.

## References

1. Ilic MZ, Handley CJ, Robinson HC, and Mok MT. Mechanism of catabolism of aggrecan by articular cartilage. Arch Biochem Biophys. 1992;294(1):115–22.

2. Loulakis P, Shrikhande A, Davis G, and Maniglia CA. N-terminal sequence of proteoglycan fragments isolated from medium of interleukin-1-treated articular-cartilage cultures. Putative site(s) of enzymic cleavage. Biochem J. 1992;284 (Pt 2)(589–93.

3. Sandy JD, Neame PJ, Boynton RE, and Flannery CR. Catabolism of aggrecan in cartilage explants. Identification of a major cleavage site within the interglobular domain. J Biol Chem. 1991;266(14):8683–5.

4. Tortorella MD, Burn TC, Pratta MA, Abbaszade I, Hollis JM, Liu R, Rosenfeld SA, Copeland RA, Decicco CP, Wynn R, et al. Purification and cloning of aggrecanase-1: a member of the ADAMTS family of proteins. Science. 1999;284(5420):1664–6.

5. Abbaszade I, Liu RQ, Yang F, Rosenfeld SA, Ross OH, Link JR, Ellis DM, Tortorella MD, Pratta MA, Hollis JM, et al. Cloning and characterization of ADAMTS11, an aggrecanase from the ADAMTS family. J Biol Chem. 1999;274(33):23443–50.

6. Little CB, Meeker CT, Golub SB, Lawlor KE, Farmer PJ, Smith SM, and Fosang AJ. Blocking aggrecanase cleavage in the aggrecan interglobular domain abrogates cartilage erosion and promotes cartilage repair. J Clin Invest. 2007;117(6):1627–36.

7. Malfait AM, Liu RQ, Ijiri K, Komiya S, and Tortorella MD. Inhibition of ADAM-TS4 and ADAM-TS5 prevents aggrecan degradation in osteoarthritic cartilage. J Biol Chem. 2002;277(25):22201–8.

8. Apte SS. Anti-ADAMTS5 monoclonal antibodies: implications for aggrecanase inhibition in osteoarthritis. Biochem J. 2016;473(1):e1–4.

9. Gilbert AM, Bikker JA, and O’Neil SV. Advances in the development of novel aggrecanase inhibitors. Expert Opin Ther Pat. 2011;21(1):1–12.

10. Miller RE, Tran PB, Ishihara S, Larkin J, and Malfait AM. Therapeutic effects of an anti-ADAMTS-5 antibody on joint damage and mechanical allodynia in a murine model of osteoarthritis. Osteoarthritis and cartilage / OARS, Osteoarthritis Research Society. 2016;24(2):299–306.

11. Fosang AJ, Neame PJ, Hardingham TE, Murphy G, and Hamilton JA. Cleavage of cartilage proteoglycan between G1 and G2 domains by stromelysins. J Biol Chem. 1991;266(24):15579–82.

12. Fosang AJ, Last K, Gardiner P, Jackson DC, and Brown L. Development of a cleavage-site-specific monoclonal antibody for detecting metalloproteinase-derived aggrecan fragments: detection of fragments in human synovial fluids. Biochem J. 1995;310 (Pt 1)(337-43.

13. Lees S, Golub SB, Last K, Zeng W, Jackson DC, Sutton P, and Fosang AJ. Bioactivity in an Aggrecan 32-mer Fragment Is Mediated via Toll-like Receptor 2. Arthritis & rheumatology. 2015;67(5):1240–9.

14. Miller RE, Belmadani A, Ishihara S, Tran PB, Ren D, Miller RJ, and Malfait AM. Damage-associated molecular patterns generated in osteoarthritis directly excite murine nociceptive neurons through Toll-like receptor 4. Arthritis & rheumatology. 2015;67(11):2933–43.

15. Kim YS, Chu Y, Han L, Li M, Li Z, Lavinka PC, Sun S, Tang Z, Park K, Caterina MJ, et al. Central terminal sensitization of TRPV1 by descending serotonergic facilitation modulates chronic pain. Neuron. 2014;81(4):873–87.

16. Miller RE, Tran PB, Das R, Ghoreishi-Haack N, Ren D, Miller RJ, and Malfait AM. CCR2 chemokine receptor signaling mediates pain in experimental osteoarthritis. Proc Natl Acad Sci U S A. 2012;109(50):20602–7.

17. Barry FP, Gaw JU, Young CN, and Neame PJ. Hyaluronan-binding region of aggrecan from pig laryngeal cartilage. Amino acid sequence, analysis of N-linked oligosaccharides and location of the keratan sulphate. Biochem J. 1992;286 (Pt 3)(761–9.

18. Brown GM, Huckerby TN, Bayliss MT, and Nieduszynski IA. Human aggrecan keratan sulfate undergoes structural changes during adolescent development. J Biol Chem. 1998;273(41):26408–14.

19. Flannery CR, Little CB, and Caterson B. Molecular cloning and sequence analysis of the aggrecan interglobular domain from porcine, equine, bovine and ovine cartilage: comparison of proteinase-susceptible regions and sites of keratan sulfate substitution. Matrix Biol. 1998;16(8):507–11.

20. Fosang AJ, Last K, Poon CJ, and Plaas AH. Keratan sulphate in the interglobular domain has a microstructure that is distinct from keratan sulphate elsewhere on pig aggrecan. Matrix Biol. 2009;28(1):53–61.

21. Liu T, Gao YJ, and Ji RR. Emerging role of Toll-like receptors in the control of pain and itch. Neurosci Bull. 2012;28(2):131–44.

22. Malfait AM, Ritchie J, Gil AS, Austin JS, Hartke J, Qin W, Tortorella MD, and Mogil JS. ADAMTS-5 deficient mice do not develop mechanical allodynia associated with osteoarthritis following medial meniscal destabilization. Osteoarthritis and cartilage / OARS, Osteoarthritis Research Society. 2010;18(4):572–80.

23. Miller RE, Tran PB, Das R, Ghoreishi-Haack N, Ren D, Miller RJ, and Malfait AM. CCR2 chemokine receptor signaling mediates pain in experimental osteoarthritis. Proceedings of the National Academy of Sciences of the United States of America. 2012;109(50):20602–7.

24. Little CB, Meeker CT, Hembry RM, Sims NA, Lawlor KE, Golub SB, Last K, and Fosang AJ. Matrix metalloproteinases are not essential for aggrecan turnover during normal skeletal growth and development. Mol Cell Biol. 2005;25(8):3388–99.

25. Miller RE, Ishihara S, Bhattacharyya B, Delaney A, Menichella DM, Miller RJ, and Malfait AM. Chemogenetic Inhibition of Pain Neurons in a Mouse Model of Osteoarthritis. Arthritis and rheumatism. In press.

26. Hayward JH, and Lee SJ. A Decade of Research on TLR2 Discovering Its Pivotal Role in Glial Activation and Neuroinflammation in Neurodegenerative Diseases. Exp Neurobiol. 2014;23(2):138–47.

27. Kim C, Rockenstein E, Spencer B, Kim HK, Adame A, Trejo M, Stafa K, Lee HJ, Lee SJ, and Masliah E. Antagonizing Neuronal Toll-like Receptor 2 Prevents Synucleinopathy by Activating Autophagy. Cell Rep. 2015;13(4):771–82.

28. Kim D, Kim MA, Cho IH, Kim MS, Lee S, Jo EK, Choi SY, Park K, Kim JS, Akira S, et al. A critical role of toll-like receptor 2 in nerve injury-induced spinal cord glial cell activation and pain hypersensitivity. J Biol Chem. 2007;282(20):14975–83.

29. Kim D, You B, Lim H, and Lee SJ. Toll-like receptor 2 contributes to chemokine gene expression and macrophage infiltration in the dorsal root ganglia after peripheral nerve injury. Mol Pain. 2011;7(74.

30. Xu ZZ, Kim YH, Bang S, Zhang Y, Berta T, Wang F, Oh SB, and Ji RR. Inhibition of mechanical allodynia in neuropathic pain by TLR5-mediated A-fiber blockade. Nat Med. 2015;21(11):1326–31.

31. Sokolove J, and Lepus CM. Role of inflammation in the pathogenesis of osteoarthritis: latest findings and interpretations. Ther Adv Musculoskelet Dis. 2013;5(2):77–94.

32. Gardiner NJ. Integrins and the extracellular matrix: key mediators of development and regeneration of the sensory nervous system. Dev Neurobiol. 2011;71(11):1054–72.

